# Flowr: Robust and efficient pipelines using a simple language-agnostic approach

**DOI:** 10.1101/029710

**Authors:** Sahil Seth, Samir Amin, Xingzhi Song, Xizeng Mao, Huandong Sun, Andrew Futreal, Jianhua Zhang

## Abstract

**Motivation:** Bioinformatics analyses have become increasingly intensive computing processes, with lowering costs and increasing numbers of samples. Each laboratory spends time creating and maintaining a set of pipelines, which may not be robust, scalable, or efficient. Further, the existence of different computing environments across institutions hinders both collabo-ration and the portability of analysis pipelines.

**Results:** Flowr is a robust and scalable framework for designing and deploying computing pipelines in an easy-to-use fashion. It implements a scatter-gather approach using computing clusters, simplifying the concept to the use of five simple terms (in submission and dependency types). Most importantly, it is flexible, such that customizing existing pipelines is easy, and since it works across several computing environments (LSF, SGE, Torque, and SLURM), it is portable.

**Availability:** http://docs.flowr.space

## Introduction

Massive advances in genomic and proteomic technologies are put-ting a high demand on bioinformatics applications for faster and more automated data processing. Since some of the steps are common and standard, there is value in creating pipelines that can be used under dif-ferent environments in various projects. In addition, several of these steps can be further broken down and parallelized to enable much faster analyses. In the past, significant efforts were made to develop tools such as Galaxy^1^ and Bpipe,^2^ ena-bling users to easily run modules and pipelines. Several other tools, such as COSMOS^3^ and BigDataScript,^4^ provide a comparatively easier syntax for building pipelines. However, all these tools require users to learn a new scripting language/syntax; thus, they present a steep learning curve. Further, such pipelines may not be portable across clusters or frameworks. Here, we present flowr, an open-source R package (http://github.com/sahilseth/flowr) that is language agnostic (in terms of inputs), robust, scalable, and portable.

## Features and Methods

One of the major challenges in creating a workflow management framework is providing essential flexibility to users without compromis-ing robustness. Flowr is language agnostic in terms of its inputs, allow-ing users to build pipelines in any language of their choice. In essence, flowr requires users to specify a set of shell commands for each step (1B), along with a simple configuration file (1C) that defines how to stitch the steps into a pipeline. Flowr provides a set of R functions for creating, reading, and checking these two input files before processing, but any other language, such as JAVA, Python, or Perl, may be used to create these simple tab-delimited text files. In addition, the configuration file (or flow definition) enables complete flexibility in specifying the computing resources, such as CPU, RAM, walltime, and queue, used in each step of the pipeline. This isolates resource specification from the actual commands, thus making the pipeline very portable across computing clusters such as LSF, Torque, SGE, SLURM, and MOAB; this is a feature unique to flowr. Further, flowr implements a scatter-gather approach, allowing a many-to-one, one-to-many, or many-to-many relationship between steps (1A and 4). Several bioinformatics pipelines can be efficiently specified using such relationships; a typical case involves processing several fastq files into a final merged binary alignment map (BAM) file. For example, several pairs of fastq files, each aligned indi-vidually using BWA^5^ (bwa aln), would be further processed as pairs using BWA (bwa sampe) to produce sam files (one for each pair). These would then be sorted, merged, and indexed using samtools.^6^ Using flowr, each step in this workflow may have different CPU, memory, and walltime usage. This simple pipeline may take up to a day on a desktop or several hours on a multicore server. Flowr efficiently scatters the steps and submits them to the cluster, man-aging dependencies, in about half an hour. The framework is robust and scalable; it creates a web of jobs (using dependencies) for the entire pipeline, submits them to the cluster, and exits. The jobs automatically start in the correct order, according to the dependency map created by flowr (example, 1E). This enables the user to submit several flows at once, in a highly scalable fashion, that will be executed depending on the resources available. Further, splitting the flow into small independent jobs enables faster processing since they fit very well in a heavily used shared computing cluster, reserving and using minimal resources. We have extensively tested flowr on several computing platforms, such as Torque, MOAB, and LSF. Using a very transparent approach, each flow is submitted as an independent container, with all commands, outputs, and logs available in a clean and structured fashion. This enables reproducibility, with the final shell scripts having all the information required to re-create the analysis. In addition, flowr creates a graph (1E) for each submission, providing a quick overview of the pipeline without reading the code. An interactive website is available for designing a new pipeline.^7^ Flowr also provides simple functions for monitoring the progress of a currently running flow, killing the whole flow, and in case of a failure, rerunning the flow from an intermediate step (2). Using a language-agnostic approach, flowr ingests the actual commands to be executed in the form of a tab-delimited file (a flowmat, 1B). Further, all the resource requirements and information regarding dependencies are isolated in a separate file (a flow definition [flowdef], 1C). The flowdef also contains information regarding the flow of steps (using the previous job column), type of submission (using the submission type column), and type of dependency on the previous jobs (using the dependency type column). Multiple commands in a module (A1-10) can be submitted in a scatter (/parallel) or serial (/sequential) fashion (1A). If a later step (B1-10) has multiple commands, such that the ith command of B depends on the ith command of A, we can describe this many-to-many relationship using a serial type dependency (1A). Further, in case of a merging step (say, C), all jobs B1-10 need to be completed, suggesting a many-to-one relationship using the dependency gather. Lastly, many steps may be initiated when this merging completes, creating a one-to-many relationship using a burst dependency (1A).

## Discussion

To our knowledge, flowr, which is explicitly based on the scatter-gather concept of data analysis pipelines, is the first open-source pipeline framework that makes use of the dependency feature of computing clusters. This feature enables flowr to intelligently submit a web of inter-dependent jobs to the computing cluster and exit, in contrast to having a daemon-type process continuously running (as in other frameworks). This minimizes overhead on the login nodes, is robust to interruptions due to accidental killing of the process, and is scalable, allowing users to submit analyses of multiple samples. Flowr follows the design once principle, enabling the user to develop robust, portable pipelines that can be run on a host of computing platforms. Further, the same pipeline can be run on a local machine, computing cluster, or cloud-based environment. With automatic logging of each step and the preservation of the exact commands run to produce the output, the system allows users to generate an easy-to-use, efficient, and reproducible analysis pipeline.

**Figure 1:**
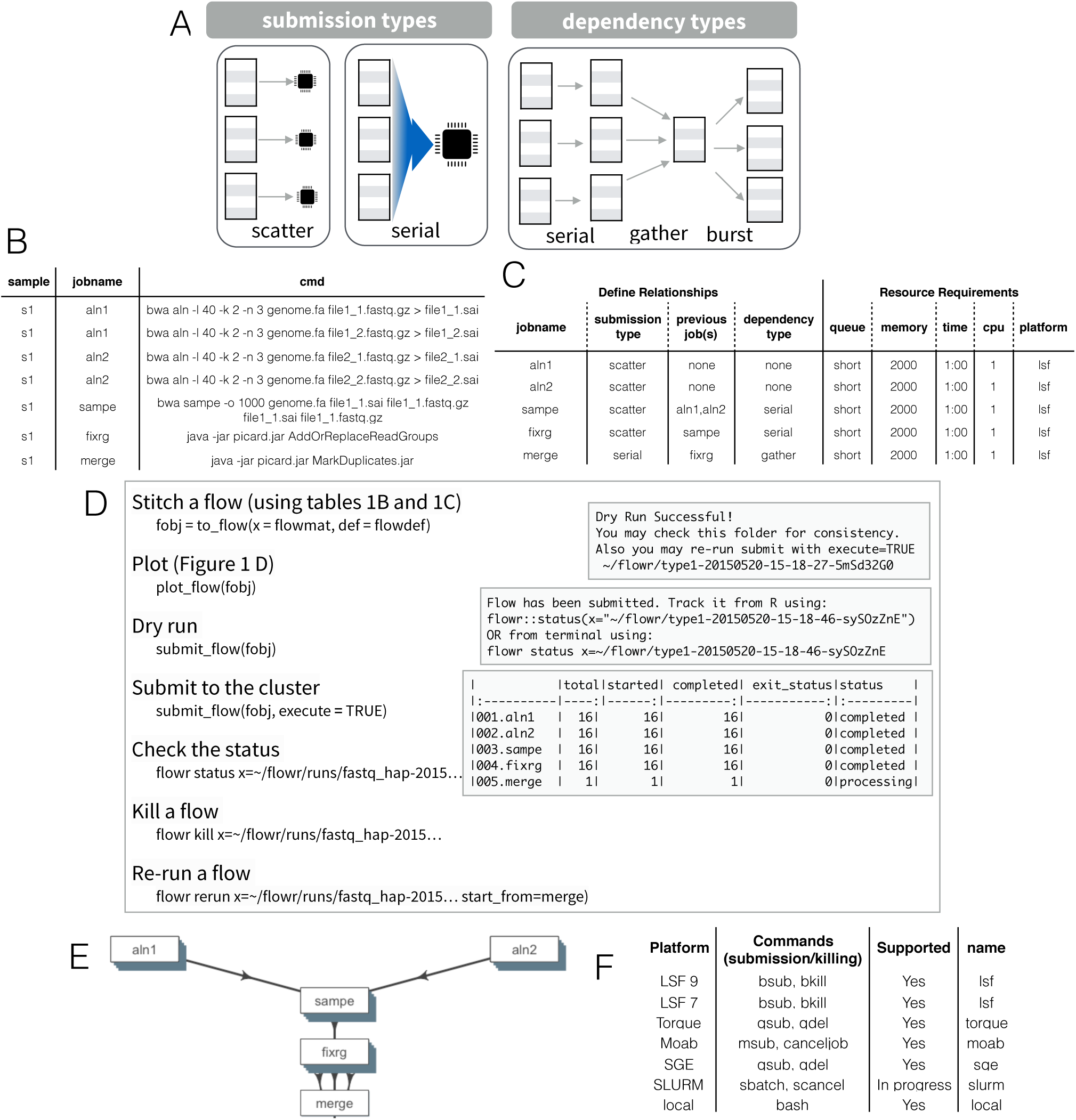
A: Among submission types, scatter submission executes jobs in parallel, while serial executes them sequentially. Gather refers to the idea that a subsequent job needs to wait for all (n) sub-processes of a previous step to complete, and serially dependent means that the ith sub-process of the current step needs to wait for the ith sub-process of a previous step. Further, burst suggests that several steps begin after a specific single step completes. We can define several complex relationships (Suppl. Table 1) using submission and dependency types. B: Flowr takes a language-agnostic approach to developing pipelines. A flow matrix (f mat) is used to describe the precise commands to run. C: A flow definition (f def) table provides an easy-to-use interface to describe various details regarding a flow, including the relationships between steps and resource requirements. Each row of the table describes one step and its relationship to previous steps, if any. Note how initial steps have none in the previous jobs and dependency type columns. D: Using a flow definition (f def) and a flow matrix (f mat), we can deploy a flow to a high-performance computing cluster. In addition, flowr provides several functions to plot the flow, monitor it, or kill and re-run it in case of issues (Suppl. 1). E: A flowchart describing procegsing the NGS workflow from fastq files to an aligned BAM file. F: Flowr supports several computing platforms out of the box, and adding support for others is quite straightforward (http://docs.flowr.space/install.html).

**Figure 2:**
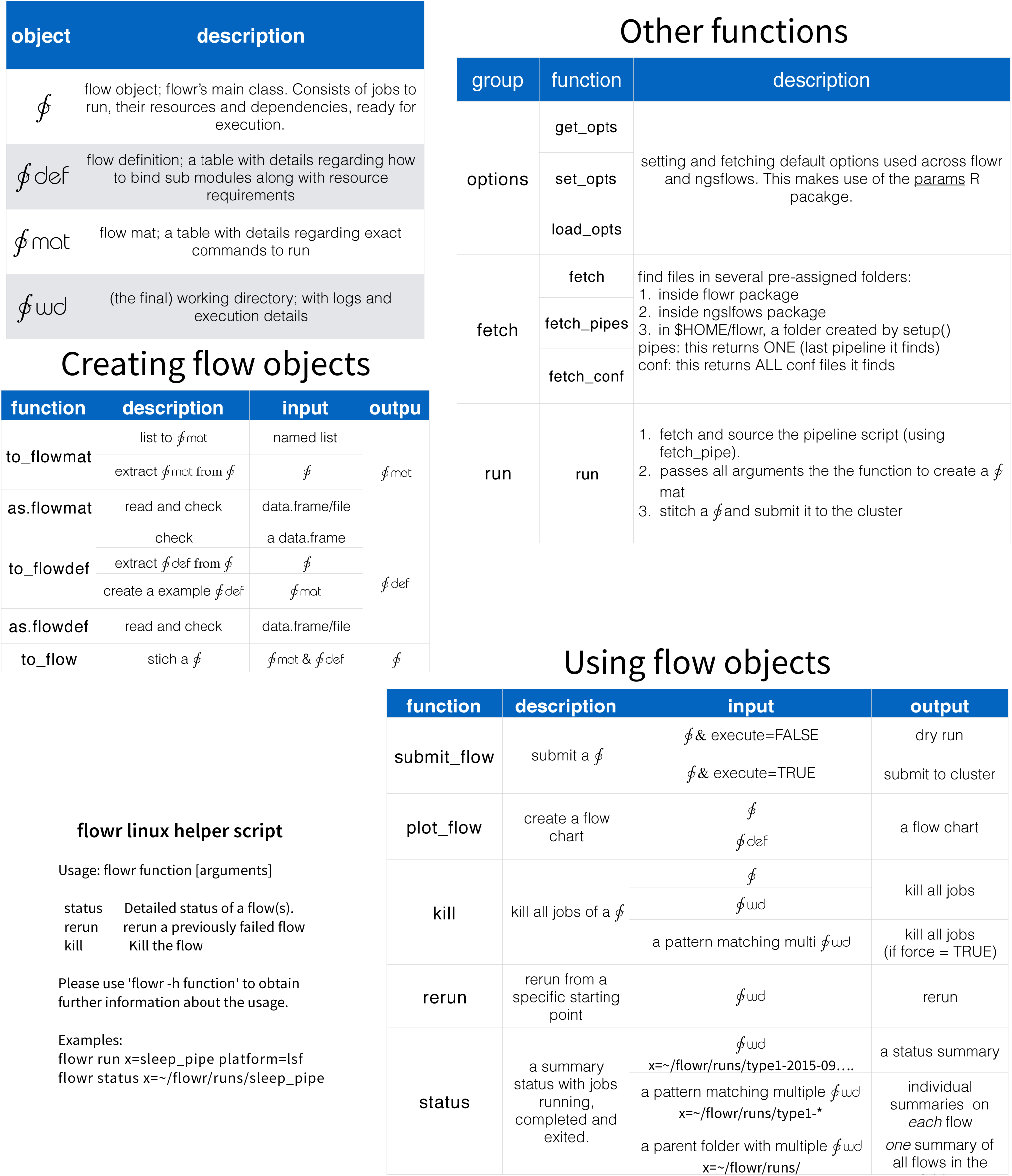
A cheat sheet describing various functions in flowr package

**Figure 3:**
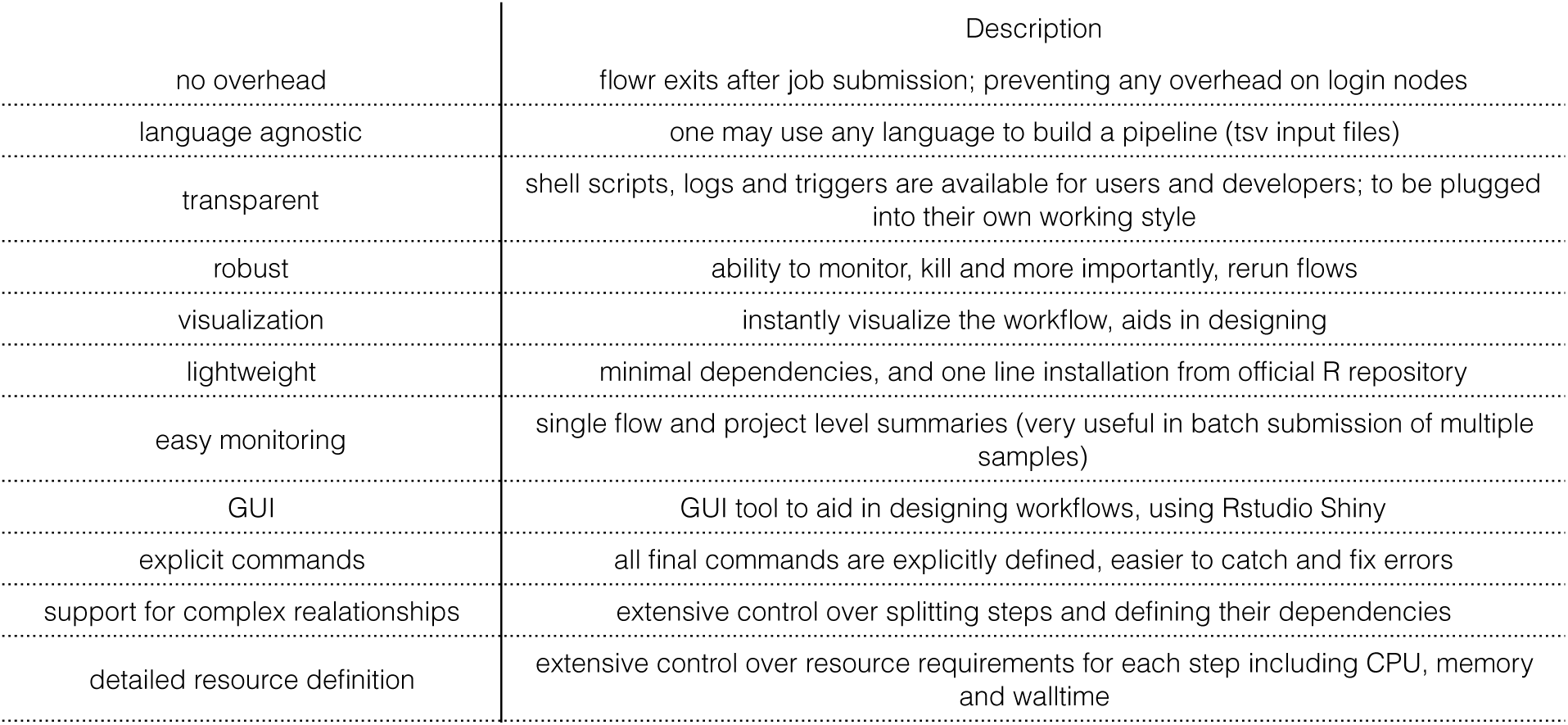
Briefly, there are several advantages of using flowr, comparing with existing workflow frameworks. Several of these stem from flowrs ability to use computing platforms dependency option.

**Figure 4:**
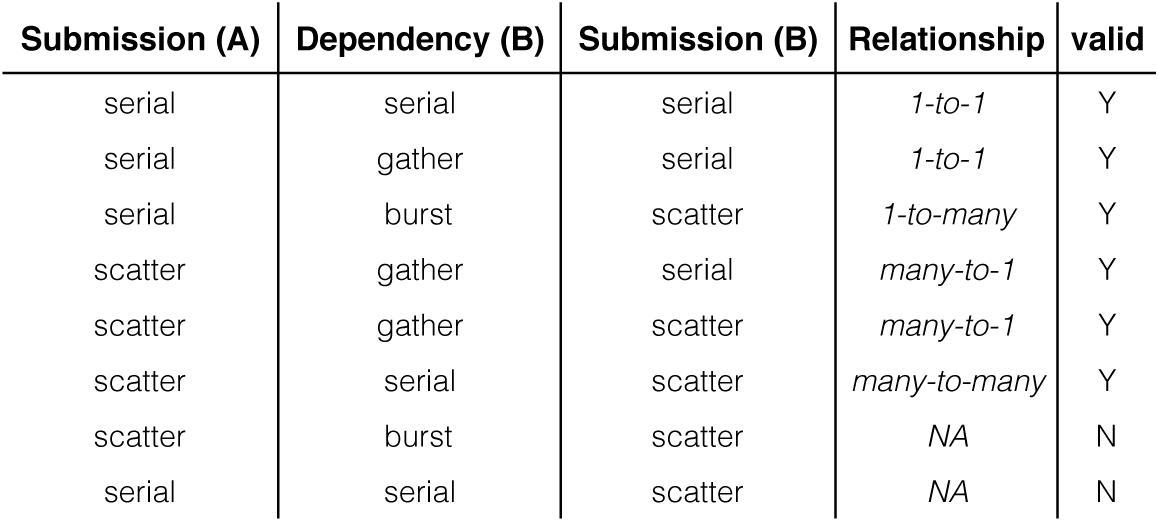
Flowr supports a functional scatter-gather approach for defining pipelines, supporting various (job) submission approaches. If a step has multiple sub-processes, a scatter approach would execute them in parallel, while serial would execute them sequentially (Figure 1B). Additionally we can define complex relationships using submission and dependency types. For example gather refers to the idea that a subsequent job needs to wait for all (n) sub-processes of a previous step to complete. Several relationships can be defined between previous (A) and subsequent jobs (B), mapping dependencies at the sub-process level. For example in many-to-many but steps (A B) have multiple sub-processing running independently in scatter mode and subprocesses in B are serially dependent means that ith subprocess of the B needs wait for the ith sub-process of a A to start.

## Acknowledgement

We thank Tapsi Seth, and members of the Verhaak, Futreal and Draetta laborato-ries for their valuable inputs We are grateful to members of the MD Anderson Research Computing team (Roger Moye, Sally Boyd, and Daniel Jackson) for their continued support. In addition, we thank Ann Sutton of the MD Anderson editorial staff, for help in editing this document.

